# Chromosome-level genome assembly and annotation of *Corallium rubrum*: a Mediterranean coral threatened by overharvesting and climate change

**DOI:** 10.1101/2024.07.13.603384

**Authors:** Jean-Baptiste Ledoux, Jessica Gomez-Garrido, Fernando Cruz, Francisco Camara Ferreira, Ana Matos, Xenia Sarropoulou, Sandra Ramirez-Calero, Didier Aurelle, Paula Lopez-Sendino, Natalie Grayson, Bradley Moore, Agostinho Antunes, Laura Aguilera, Marta Gut, Judit Salces-Ortiz, Rosa Fernández, Cristina Linares, Joaquim Garrabou, Tyler Alioto

## Abstract

Reference genomes are key resources in biodiversity conservation. Yet, sequencing efforts are not evenly distributed in the tree of life questioning our true ability to enlighten conservation with genomic data. Good quality reference genomes remain scarce in octocorals while these species are highly relevant target for conservation. Here, we present the first annotated reference genome in the red coral, *Corallium rubrum* (Linnaeus, 1758), a habitat-forming octocoral from the Mediterranean and neighboring Atlantic, impacted by overharvesting and anthropogenic warming-induced mass mortality events. Combining long reads from Oxford Nanopore Technologies (ONT), Illumina paired-end reads for improving the base accuracy of the ONT-based genome assembly and Arima Hi-C contact data to place the sequences into chromosomes, we assembled a genome of 475 Mb (21 chromosomes, 326 scaffolds) with contig and scaffold N50 of 1.6 Mb and 16.2 Mb, respectively. Fifty percent of the sequence (L50) was contained in eight superscaffolds. The consensus quality (QV) of the final assembly was 42 and the gene completeness reported by BUSCO was 74% (metazoa_odb10 database). We annotated 39,114 protein-coding genes and 32,678 non-coding transcripts. This annotated chromosome-level genome assembly, one of the first in octocorals, is currently used in a project based on whole genome re-sequencing dedicated to the conservation and management of *C. rubrum*.

**Significance Statement:** The Mediterranean red coral, *Corallium rubrum*, is critically impacted by overharvesting and by mass mortality events linked to marine heat waves. Accordingly, *C. rubrum* is increasingly receiving conservation efforts. Previous population genetics studies based on microsatellites contributed to improving our knowledge of the species ecology. Yet, crucial questions regarding, admixture among lineages, demographic history, effective population sizes and local adaptation, are still open owing to a lack of genomic resources. Here, we present the first chromosome-level genome assembly for the species with high contiguity, good completeness and protein-coding genes and repeat sequence annotations. This genome, one of the first in octocorals, will pave the way for the integration of population genomics data into ongoing interdisciplinary conservation efforts dedicated to *C. rubrum*.

## Introduction

Recent improvements in sequencing technologies and bioinformatics are upgrading the potential inputs of population genetics in conservation biology (Formenti et al. 2022). Accordingly, the number of initiatives to produce high-quality reference genomes has been increasing in the last five years (e.g. Catalan initiative for the Earth Biogenome Project https://www.biogenoma.cat).

Yet, these efforts are still mostly focused on a few taxa (e.g. Vertebrate Genome Project) and reference genomes for non-model Metazoans are still scarce (but see Ledoux et al. 2020). This bias in the sequencing efforts is detrimental to biodiversity conservation owing to the ecological roles of many of those underrepresented taxa.

Beyond an interesting phylogenetic position as a sister taxa of Hexacorallia, Octocorallia is a diverse group (>3,500 species) of ecologically key organisms (e.g. Gomez-Gras et al. 2021) found from shallow tropical to deep and polar seas. Some of these species are critically impacted by global change including extreme climatic events (e.g. Estaque et al. 2023). To date, only a bunch of genomes (<1% of species diversity, see Ahuja et al. 2024) are available limiting the integration of genomics data into ongoing conservation efforts.

The red coral, *Corallium rubrum*, is a habitat-forming octocoral (Figure 1) with a central structural role in benthic communities from the Mediterranean and the neighboring Atlantic (Zibrowius et al. 1984, Laborel & Vacelet 1961). This iconic species with high cultural and economic value is critically impacted by two anthropogenic pressures. First, as a “precious coral”, it has been harvested for jewelry since ancient times and owing to its market value (>1,000 €/kg), the species has been overharvested and intensively poached (Ledoux et al. 2016). Second, *C. rubrum* has been recurrently impacted in the last twenty years by mass mortalities, linked to recurrent marine heatwaves, across thousands of kilometers of coastal habitats (Garrabou et al. 2022). The species with slow population dynamics (Montero-Serra et al. 2018) and restricted connectivity (Ledoux et al. 2010a; Horaud et al. 2023) is characterized by a low resilience capacity (Linares et al. 2012). The combination of overharvesting and mass mortality events is driving steep demographic declines, questioning the evolutionary trajectory of the species (Montero-Serra et al. 2019).

**Figure 1:**
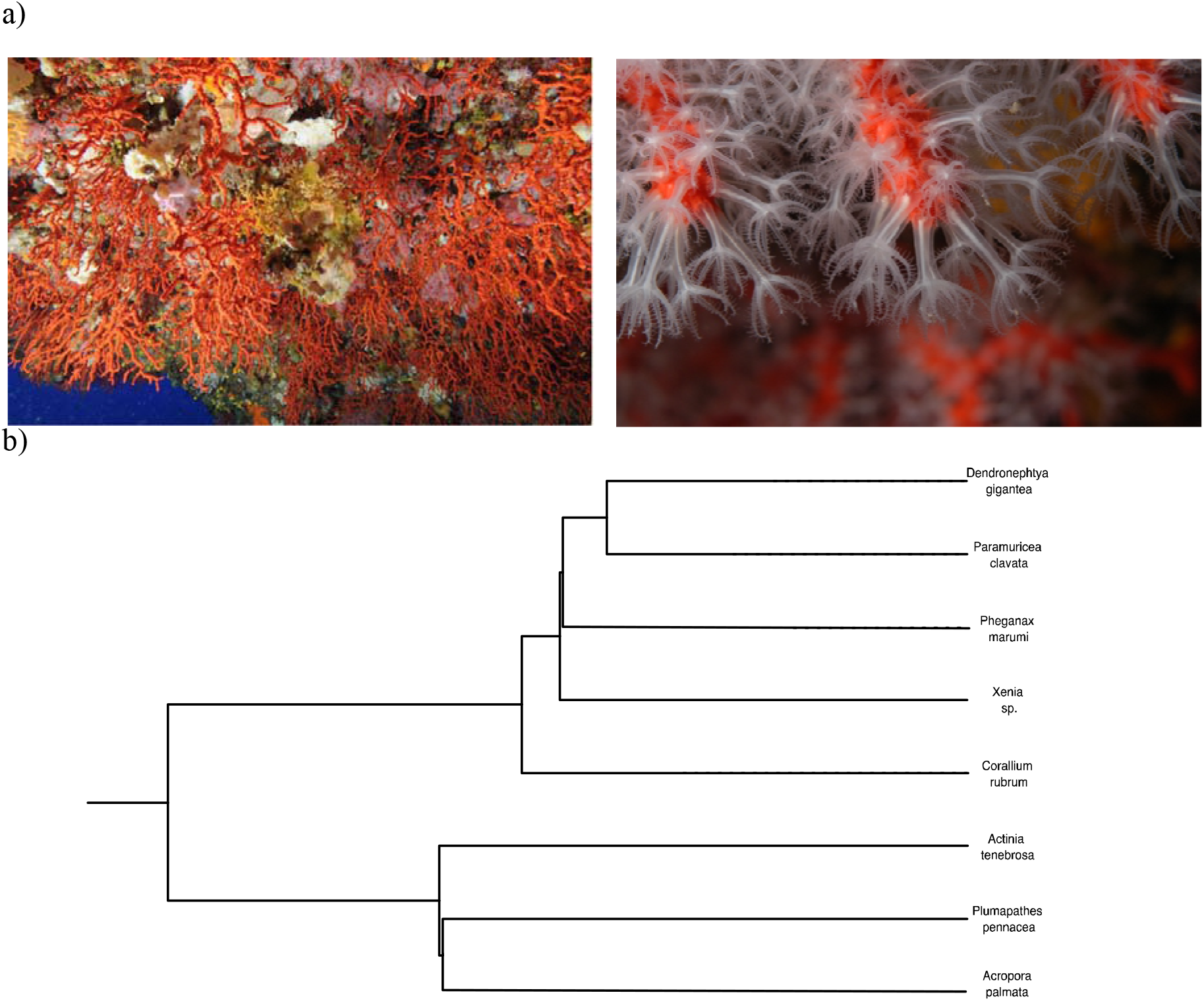
a) Coralligenous habitat dominated by the red coral, *Corallium rubrum* (left panel). Close up from apical tips of *C. rubrum* showing the polyps (white) and coenenchyma covering the red calcareous skeleton used in jewelry since Ancient time (right panel). b) Phylogenetic relationships among different anthozoans species including five octocorals (*Dendronephtya gigantea, Paramuricea clavata, Pheganax marumi, Xenia sp*., *Corallium rubrum*) and three hexacorals (*Actinia tenebrosa, Plumathes pennacea, Acropora palmata*) for which good quality assemblies are available. species with good quality genome assemblies. The tree is based on 244 single copy orthologous genes identified with BUSCO.

In this context, *C. rubrum* is receiving conservation attention from scientists and biodiversity managers (included in Barcelona Convention, EU Habitat Directive and listed as “endangered” by IUCN [Otero et al. 2017]). Yet, major knowledge gaps in relation to genome diversity, effective population size and adaptation to the local environment remain and should be filled to improve existing conservation policies. As a part of the Catalan Initiative for the Earth BioGenome Project (CBP), we assembled and annotated the first chromosome-level reference genome in *C. rubrum*. This reference genome will support a conservation genomics project funded by the Biodiversity Genomics Europe (https://biodiversitygenomics.eu) and based on whole genome re-sequencing. This project will infer demographic history and contemporary processes shaping the intraspecific genetic patterns with direct applications for red coral conservation and management.

## Material and Methods

### Collection and preparation of biological material

The apical tip (5 cm) of one colony from the Cap Castell (42.082610, 3.201981) population in Catalunya (Spain) was sampled at 18 m depth and immediately transported in coolers to the Aquarium Experimental Zone (ZAE) of the Institut de Ciències del Mar (ICM-CSIC, Barcelona, Spain). The sample was flash frozen using liquid nitrogen and conserved at -80°C until DNA extractions. The same individual was used for short (Illumina) and long-read (Oxford Nanopore Technology) sequencing. For Hi-C sequencing, one individual colony was sampled from Meda Petita population at 12m depth (42.043652; 3.226719), Medes Islands, Spain.

### DNA extraction and Illumina Whole Genome Sequencing

High Molecular Weight gDNA was extracted from the coenenchyme (external tissue containing the polyps) using the MagAttract HMW DNA kit (Qiagen) at the Centre Nacional d’Analisi Genomica (CNAG, https://www.cnag.eu). The HMW gDNA eluate was quantified using the Qubit DNA BR Assay kit (Thermo Fisher Scientific), and its purity was assessed using Nanodrop 2000 (Thermo Fisher Scientific). The extractions integrity was analyzed in an agarose gel (1%) in a pulsed field gel electrophoresis system (Sage Science). The HMW gDNA sample was stored at 4°C. Whole genome sequencing library preparation was performed using the KAPA HyperPrep kit (Roche), following the manufacturer’s instructions. The libraries were sequenced on the NovaSeq 6000 (Illumina) with a read length of 2×151bp, following the manufacturer’s protocol for dual indexing. Image analysis, base calling, and quality scoring of the run were executed using the manufacturer’s Real Time Analysis (RTA 3.4.4) software.

### Long-Read Whole genome library preparation and sequencing

The sequencing libraries were prepared using the 1D Sequencing kit SQK-LSK110 from Oxford Nanopore Technologies (ONT). Briefly, 4.0 μg of the DNA was DNA-repaired and DNA-end-repaired using NEBNext FFPE DNA Repair Mix (NEB) and the NEBNext UltraII End Repair/dA-Tailing Module (NEB) followed by the sequencing adaptors ligation. The ligation product was purified by 0.4X AMPure XP beads (Agencourt, Beckman Coulter), and eluted in elution buffer.

The sequencing runs were performed on PromethIon 24 (ONT) using a flow cell R9.4.1 FLO-PRO 002 (ONT) and the sequencing data was collected for 110 hours. The quality parameters of the sequencing runs were monitored by the MinKNOW platform version 21.11.7 in real time and base called with Guppy version 5.1.13.

### Chromatin conformation capture sample preparation and sequencing

Tissue was carefully scraped from a living individual collected at Medas Petit. Chromatin conformation capture sequencing (Hi-C) libraries were prepared using the Hi-C High-coverage kit (Arima Genomics) in the Metazoa Phylogenomics Lab (Institute of Evolutionary Biology (CSIC-UPF)). Sample concentration was assessed by Qubit DNA HS Assay kit (Thermo Fisher Scientific) and library preparation was carried out using the ACCEL-NGS®□ 2S PLUS DNA LIBRARY KIT (Swift Bioscience) and using the 2S Set A single indexes (Swift Bioscience). Library amplification was carried out with the KAPA HiFi DNA polymerase (Roche). The amplified libraries were sequenced on the NovaSeq 6000 (Illumina) at CNAG.

### RNA extraction and RNA Sequencing

RNA sequencing data were obtained from a parallel project characterizing the transcriptomic response of *C. rubrum* to heat stress (Ramirez et al. in prep). RNA was extracted from the coenenchyme of 36 different samples combining TRIzol reagent (Invitrogen) for tissue lysis and homogenization and RNA easy kit (Qiagen) for RNA isolation and purification. Eluted RNA was stored at -80°C until shipment to CNAG. Total RNA quantification was assessed using the Qubit RNA BR Assay kit (Thermo Fisher Scientific), and the RNA integrity was estimated using the RNA 6000 Nano Bioanalyzer 2100 Assay (Agilent). To prepare the RNA-Seq libraries, the KAPA Stranded mRNA-Seq Illumina Platforms Kit (Roche) was used with 500 ng of total RNA. Library quality was assessed on an Agilent 2100 Bioanalyzer using the DNA 7500 assay. The libraries were sequenced on the NovaSeq 6000 (Illumina) as above for the WGS library.

### Genome assembly

We used the pipeline CLAWS v2.1 (Gomez-Garrido, 2023) to perform this genome assembly combining ONT long reads, Illumina paired-end reads and Arima Hi-C contact data. A flowchart with the genome assembly process is shown in Supplementary file (Figure S1).

Prior to assembly, adaptors present in the Illumina data were trimmed with TrimGalore (https://github.com/FelixKrueger/TrimGalore). A k-mer database was subsequently built with Meryl (https://github.com/marbl/meryl). The k-mer histogram generated by Meryl was used as input to Genomescope2 (Ranallo-Benavidez et al. 2020) to estimate haploid genome size, heterozygosity and repeat content (Supplementary file Figure S2). The ONT data were filtered with Filtlong (https://github.com/rrwick/Filtlong; *--minlen* 1000 *--min_mean_q* 80 *--target_bases* 25000000000) prior to the assembly to remove short and low-quality reads.

The filtered ONT data was assembled with Nextdenovo v2.4.0 (Hu et al. 2024). To improve the base accuracy, the assembly was polished with HyPo (Kundu et al. 2019) using both Illumina and ONT data. Finally, the polished assembly was purged with *purge_dups* (Guan et al. 2020) to remove alternate haplotypes and other artificially duplicated repetitive regions.

The Blobtoolkit (Challice et al. 2020) pipeline was run, using the NCBI nucleotide database (updated in February 2023) and several BUSCO odb10 databases (metazoa, eukaryota, fungi and bacteria). A total of 135 contigs (corresponding to 70.2 Mb of sequences) belonging to non-Cnidaria phyla were removed from the assembly at this step (see blobplot Figure S3).

The decontaminated assembly was scaffolded using the Hi-C data with YAHS (Zhou et al. 2022). Manual curation of the resulting assembly was performed with PretextView (https://github.com/wtsi-hpag/PretextView). A total of 124 edits were made (183 interventions per gigabase), of which 29 were breaks and 58, joins. The rest corresponded to 36 unlocalized sequences and one haplotig. A total of 21 autosomes were assembled and no sex chromosomes were identified.

A snailplot was produced on the final assembly with Blobtoolkit (Figure 2).

**Figure 2:**
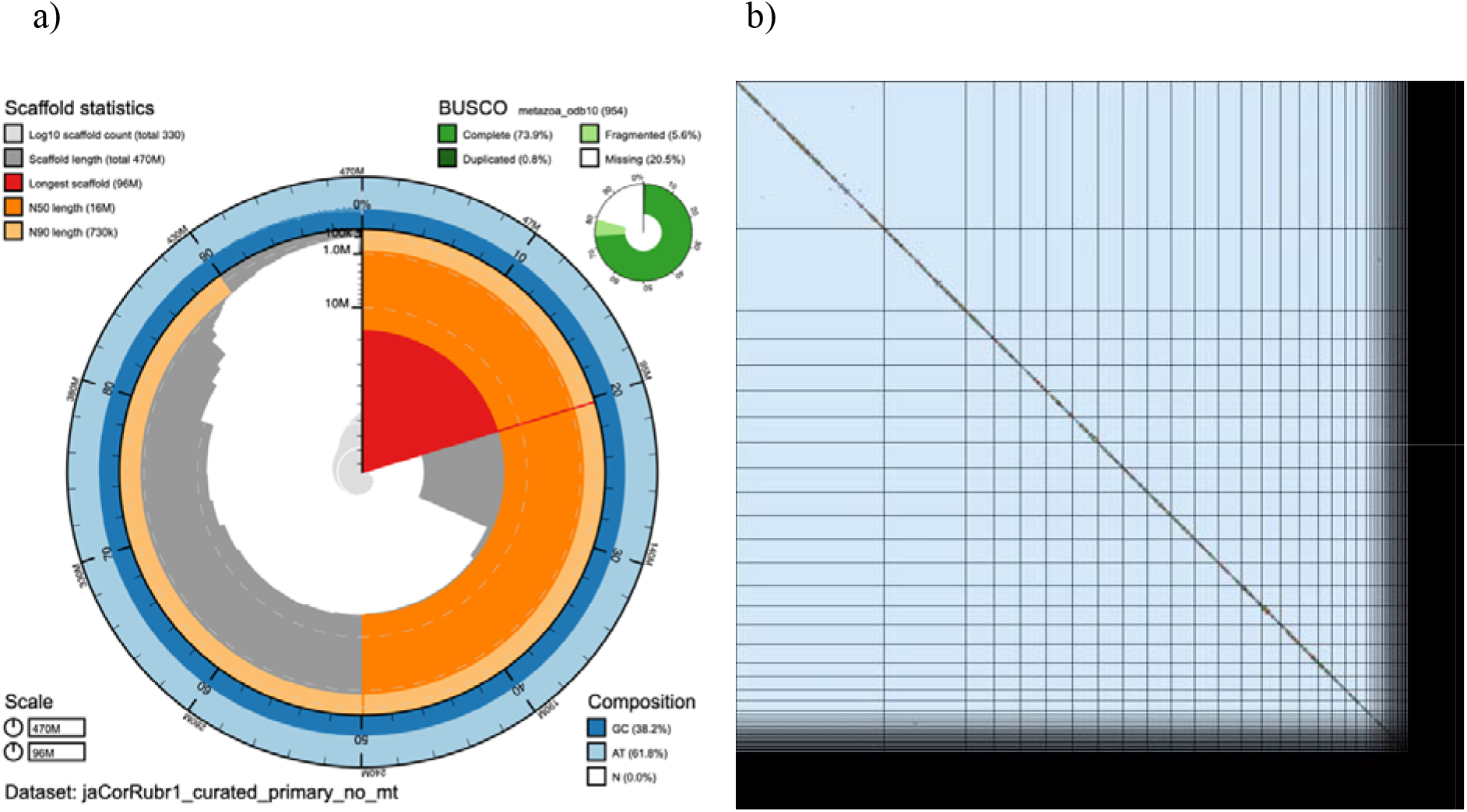
a) BlobToolKit Snailplot showing different assembly metrics. The main plot is divided into 1,000 size-ordered bins around the circumference with each bin representing 0.1% of the 474.689.186 bp assembly. The distribution of scaffold lengths is shown in dark grey with the plot radius scaled to the longest scaffold present in the assembly (96.441.827 bp, shown in red). Orange and pale-orange arcs show the N50 and N90 scaffold lengths (16.290.029 and 728.786 bp), respectively. The pale grey spiral shows the cumulative scaffold count on a log scale with white scale lines showing successive orders of magnitude. The blue and pale-blue area around the outside of the plot shows the distribution of GC, AT and N percentages in the same bins as the inner plot. A summary of complete, fragmented, duplicated and missing BUSCO genes in the metazoa_odb10 set is shown in the top right. b) Chromatin contact map generated from Arima2 Hi-C data shows the 21 chromosomes (2*n* = 42) that represent 88.8% of the assembled *C. rubrum* genome.

### Genome annotation

The genome annotation was obtained by running the CNAG structural genome annotation pipeline (https://github.com/cnag-aat/Annotation_AAT) that uses a combination of transcript alignments, protein alignments and *ab initio* gene predictions (Supplementary file Figure S4). Repeats present in the genome assembly were annotated with RedMask. To avoid masking certain repetitive protein families present in the genome, we performed a BLAST (Altschul et al. 1990) search of the RedMask-produced library against Swissprot/Uniprot (February 2023). Those repeats with significant hits (evalue <10^−6^) against proteins were removed from the final repeat library and BedTools v2.31.1 (Quinlan & Hall 2010) was run to produce the masked version of the genome.

After sequencing, adaptors were removed from the reads corresponding to the 36 samples with TrimGalore. Reads were aligned to the genome with STAR v-2.7.2a (Dobin et al. 2013). Transcript models were subsequently generated using Stringtie v2.2.1 (Pertea et al. 2015) on each BAM file and then all the models produced were combined using TACO v0.7.3 (Niknafs et al. 2017). High-quality junctions used during the annotation process were obtained by running ESPRESSO v1.3.0 (Gao et al. 2023) after mapping with STAR. Finally, PASA assemblies were produced with PASA v2.5.2 (Haas et al. 2015). The *TransDecoder* program was run on the PASA assemblies to detect the presence of coding regions in the transcripts. Additionally, the complete proteomes of *Stylopora pistillata, Pocillopora damicornis* and *Paramuricea clavata* were downloaded from Swissprot/Uniprot (February 2023) and aligned to the *C. rubrum* genome using Miniprot v0.6 (Li 2023). *Ab initio* gene predictions were performed on the repeat-masked assembly with three different programs: GeneID v1.4 (Alioto et al. 2018), Augustus v3.5.0 (Stanke et al. 2006) and Genemark-ET v7.71 (Lomsadze et al. 2014) with and without incorporating evidence from the RNAseq data. Geneid and Augustus were specifically trained for this species using a set of 1000 gene candidates obtained from the longest Transdecoder complete models that had a significant (evalue <10^−6^) BLAST hit against Swissprot/Uniprot. Genemark was run in a self-training mode and it was not specifically trained with this set of gene candidates.

Finally, all the data were combined into consensus CDS models using EvidenceModeler-2.1 (Haas et al. 2015). Additionally, UTRs and alternative splicing forms were annotated via two rounds of PASA annotation updates. To functionally annotate the proteins of the annotation, we run the Pannzer’s online server (Törönen & Holm 2020). Orthofinder (Emms & Kelly 2019) was run to obtain the orthologs between *C. rubrum* and the previously downloaded proteins for *P. clavata, P. damicornis* and *S. pistillata*. The proteins that had not originally been annotated by Pannzer but for which an ortholog was found, inherited the functional tags of their other paralogs in the *C. rubrum* annotation or, if absent, they hierarchically obtained the annotation of their orthologs in *P. clavata, P. damicornis* or *S. pistillata*.

The annotation of ncRNAs was obtained by running the following steps on the repeat-masked version of the genome assembly. First, cmsearch v1.1 (Cui et al. 2016) that is part of the Infernal package (Nawrocki et al. 2013) was run against the RFAM database of RNA families v12.0. Additionally, tRNAscan-SE v2.11 (Chan & Lowe 2017) was run to identify the transfer RNA genes present in the genome assembly. Identification of lncRNAs was done by first filtering the set of PASA-assemblies that had not been included in the annotation of protein-coding genes to retain those longer than 200bp and not covered more than 80% by a small ncRNA. The resulting transcripts were clustered into genes using shared splice sites or significant sequence overlap as criteria for designation as the same gene.

## Results and Discussion

### Genome assembly

Results obtained with Genomescope2 (Figure S2) suggest a genome-size of around 500 Mb and 1.2% heterozygosity rate. The base assembly obtained with NextDenovo v2.4.1 comprised a total assembly span of 568 Mb (876 contigs) and the final chromosome-level assembly comprised 475 Mb (21 chromosomes, 326 scaffolds) (Table S1). The contig and scaffold N50 of the final assembly are 1.6 Mb and 16.2 Mb, respectively, and fifty percent of the sequence (L50) is placed in eight superscaffolds. BUSCO (Manni et al. 2021) and Merqury (Rhie et al. 2020) were run to estimate the accuracy and completeness of the genome assembly. The consensus quality (QV) of the final assembly was estimated by Merqury as 42 and the gene completeness reported by BUSCO v5 was 74% using the *metazoa_odb10* database (Figure 2; Table S1).

### Genome annotation

We annotated a total of 39,114 protein-coding genes that produce 44,624 transcripts (1.14 transcripts per gene) and encode for 43,533 unique protein products. We were able to assign functional labels to 45% of the annotated proteins. The annotated transcripts contain 5.09 exons on average, with 63% of them being multi-exonic (Table S2). In addition, 32,678 non-coding transcripts were annotated, of which 24,752 and 7,926 are long and short non-coding RNA genes, respectively.

The reference genome presented here is the backbone of an ongoing population genomics project dedicated to the conservation and management of *C. rubrum*. This chromosome-level assembly, one of the first in octocorals and the first in within the order Scleralcyonacea, contributes to reduce the current taxonomic bias in the generation of high-quality genome resources.

## Data Availability

Data and genome assembly presented in this article are available from CNAG (https://denovo.cnag.cat/) and ENA (Project GCA_964035015.1 ; https://www.ebi.ac.uk/ena/browser/view/GCA_964035015.1).

## Acknowledgement

This project was supported by the first call for sequencing reference genomes from the Catalan Initiative for the Earth Biogenome Project. JBL was supported by the strategic funding UIDB/04423/2020, UIDP/04423/2020 and 2021.00855.CEECIND through national funds provided by FCT -Fundaço para a Ciência e a Tecnologia. Institutional support to CNAG was from the Spanish Ministry of Science and Innovation through the Instituto de Salud Carlos III and Generalitat de Catalunya through the Departament de Salut and the Departament de Recerca i Universitats. The project leading to this publication has received funding from European FEDER Fund under project 1166-39417. The project leading to this publication has received funding from Excellence Initiative of Aix-Marseille University - A*MIDEX, a French “Investissements d’Avenir” programme. RF acknowledges support from the following sources of funding: Ramón y Cajal fellowship (grant agreement no. RYC2017-22492 funded by MCIN/AEI /10.13039/501100011033 and ESF ‘Investing in your future’), the Agencia Estatal de Investigación (project PID2019-108824GA-I00 funded by MCIN/AEI/10.13039/501100011033), the European Research Council (this project has received funding from the European Research Council (ERC) under the European’s Union’s Horizon 2020 research and innovation programme (grant agreement no. 948281)) and the Secretaria d’Universitats i Recerca del Departament d’Economia i Coneixement de la Generalitat de Catalunya (AGAUR 2021-SGR00420). JG acknowledges the funding of the Spanish government through the ‘Severo Ochoa Centre of Excellence’ accreditation (CEX2019-000928-S). CL gratefully acknowledges the financial support by ICREA under the ICREA Academia programme. LFF was supported by an FI SDUR grant (2020 FISDU 00482) from the ‘Generalitat de Catalunya’. JG, CL, PLS, SRC and JBL are part of the Marine Conservation research group –MedRecover (2021 SGR 01073) from the ‘Generalitat de Catalunya’.

## References

Ahuja, M. et al. 2024 Giants among Cnidaria: Large Nuclear Genomes and Rearranged Mitochondrial Genomes in Siphonophores, Genome Biology and Evolution, Volume 16, Issue 3, 10.1093/gbe/evae048

Alioto, T., Blanco, E., Parra, G., and Guigó, R. 2018, Using geneid to Identify Genes. Curr Protoc Bioinformatics, 64, e56.

Altschul, S. F., Gish, W., Miller, W., Myers, E. W., and Lipman, D. J. 1990, Basic local alignment search tool. J Mol Biol, 215, 403–10.

Challis, R., Richards, E., Rajan, J., Cochrane, G., and Blaxter, M. 2020, BlobToolKit – Interactive Quality Assessment of Genome Assemblies. G3 Genes|Genomes|Genetics, 10, 1361–74.

Chan, P. P., and Lowe, T. M. 2019, tRNAscan-SE: Searching for tRNA Genes in Genomic Sequences. Methods Mol Biol, 1962, 1–14.

Cui, X., et al. 2016, CMsearch: simultaneous exploration of protein sequence space and structure space improves not only protein homology detection but also protein structure prediction. Bioinformatics, 32, i332–40.

Dobin, A., et al. 2013, STAR: ultrafast universal RNA-seq aligner. Bioinformatics, 29, 15–21.

Emms, D. M., and Kelly, S. 2019, OrthoFinder: phylogenetic orthology inference for comparative genomics. Genome Biol, 20, 238.

Estaque, T., et al. 2022. (2023), Marine heatwaves on the rise: One of the strongest ever observed mass mortality event in temperate gorgonians. Glob Change Biol, 29: 6159–6162. 10.1111/gcb.16931

Formenti, G, et al. 2022. The era of reference genomes in conservation genomics. Trends Ecol. Evol. 37, 197–202.

Gao, Y., et al. 2023, ESPRESSO: Robust discovery and quantification of transcript isoforms from error-prone long-read RNA-seq data. Sci. Adv., 9, eabq5072.

Garrabou, J., et al. (2022). Marine heatwaves drive recurrent mass mortalities in the Mediterranean Sea. Glob Ch Bio, 28, 5708–5725.

Gomez-Garrido, J. 2023, CLAWS (CNAG’s long-read assembly workflow for Snakemake).

Gómez-Gras, D., et al. (2021), Climate change transforms the functional identity of Mediterranean coralligenous assemblages. Ecology Letters, 24: 1038–1051. 10.1111/ele.13718

Guan, D., et al. 2020, Identifying and removing haplotypic duplication in primary genome assemblies. Bioinformatics, 36, 2896–8.

Haas, B. J., et al. 2008, Automated eukaryotic gene structure annotation using EVidenceModeler and the Program to Assemble Spliced Alignments. Genome Biol, 9, R7.

Hu, J., Wang, Z., Sun, Z., et al. 2024, NextDenovo: an efficient error correction and accurate assembly tool for noisy long reads. Genome Biol, 25, 107.

Kundu, R., Casey, J., and Sung, W.-K. 2019, HyPo: Super Fast & Accurate Polisher for Long Read Genome Assemblies. preprint, Bioinformatics.

Laborel J, Vacelet J (1961) Répartition bionomique du Corallium rubrum LMCK dans les grottes et falaises sous-marines. Rapports et Procès-Verbaux des Réunions de la Commission Internationale pour l’Exploration Scientifique de la Mer Méditerranée, 16, 464–469.

Ledoux J.-B., et al. (2010) Genetic survey of shallow populations of the Mediterranean red coral (Corallium rubrum (Linnaeus, 1758)): new insights into evolutionary processes shaping current nuclear diversity and implications for conservation. Mol. Ecol., 19, 675–690.

Ledoux JB., et al. 2016 Molecular forensics into the sea: how molecular markers can help to struggle against poaching and illegal trade in precious corals? pp. 729-746. in Medusa and her children. Cnidarian evolution, through global climate change effects (SPRINGER eds).

Ledoux J.-B., et al. 2020. The genome sequence of the octocoral Paramuricea clavata – a key resource to study the impact of climate change in the Mediterranean. G3 Gene Genome Genetics 10, 2941-2952. 10.1101/849158.

Li, H. 2023, Protein-to-genome alignment with miniprot. Bioinformatics, 39, btad014.

Linares, C., et al. 2012, Assessing the Effectiveness of Marine Reserves on Unsustainably Harvested Long-Lived Sessile Invertebrates. Conservation Biology, 26: 88–96. 10.1111/j.1523-1739.2011.01795.x

Lomsadze, A., Burns, P. D., and Borodovsky, M. 2014, Integration of mapped RNA-Seq reads into automatic training of eukaryotic gene finding algorithm. Nucleic Acids Res, 42, e119.

Manni, M., Berkeley, M. R., Seppey, M., Simão, F. A., and Zdobnov, E. M. 2021, BUSCO Update: Novel and Streamlined Workflows along with Broader and Deeper Phylogenetic Coverage for Scoring of Eukaryotic, Prokaryotic, and Viral Genomes. Molecular Biology and Evolution, 38, 4647–54.

Montero-Serra I. Linares C., Doak D.F., Ledoux J.-B., Garrabou J. (2018) Strong linkages between depth, longevity and demographic stability across marine sessile species. Proceedings of the Royal Society B 285, 1873. DOI: 10.1098/rspb.2017.2688

Montero-Serra I., Garrabou J., Doak DF., Ledoux J.-B., Linares C. (2019) Marine protected areas enhance structural complexity but do not buffer the consequences of ocean warming for an overexploited precious coral. Journal of Applied Ecology DOI: 10.1111/1365-2664.13321

Nawrocki, E. P., and Eddy, S. R. 2013, Infernal 1.1: 100-fold faster RNA homology searches. Bioinformatics, 29, 2933–5.

Niknafs, Y. S., Pandian, B., Iyer, H. K., Chinnaiyan, A. M., and Iyer, M. K. 2017, TACO produces robust multisample transcriptome assemblies from RNA-seq. Nat Methods, 14, 68–70.

Otero, M. et al. (2017). Overview of the conservation status of Mediterranean anthozoa. Overview of the Conservation Status of Mediterranean Anthozoa.

Pertea, M., et al. 2015, StringTie enables improved reconstruction of a transcriptome from RNA-seq reads. Nat Biotechnol, 33, 290–5.

Quinlan, A. R., and Hall, I. M. 2010, BEDTools: a flexible suite of utilities for comparing genomic features. Bioinformatics, 26, 841–2.

Ranallo-Benavidez, T. R., Jaron, K. S., and Schatz, M. C. 2020, GenomeScope 2.0 and Smudgeplot for reference-free profiling of polyploid genomes. Nat Commun, 11, 1432.

Rhie, A., Walenz, B. P., Koren, S., and Phillippy, A. M. 2020, Merqury: reference-free quality, completeness, and phasing assessment for genome assemblies. Genome Biology, 21, 245.

Stanke, M., Schöffmann, O., Morgenstern, B., and Waack, S. 2006, Gene prediction in eukaryotes with a generalized hidden Markov model that uses hints from external sources. BMC Bioinformatics, 7, 62.

Törönen, P., and Holm, L. 2022, PANNZER —A practical tool for protein function prediction. Protein Science, 31, 118–28.

Wick, R. FiltLong.

Zibrowius H, Monteiro-Marques V, Grasshoff M (1984) La répartition du Corallium rubrum dans l’Atlantique (Cnidaria: Anthozoa: Gorgonaria). Téthys, 11, 163–170.

Zhou, C., McCarthy, S. A., and Durbin, R. 2022, YaHS: yet another Hi-C scaffolding tool. bioRxiv, 2022.06.09.495093.

