## supplementary file for "Chromosome-level genome assembly and annotation of *Corallium rubrum*: a Mediterranean coral threatened by overharvesting and climate change"

Supplementary information

In this Supplementary Information, we describe:

- The workflow of the genome assembly process (Figure S1)
- The linear plot from Genomescope2 (Figure S2)
- The blobplot of base coverage function of GC content (Figure S3)
- The workflow of the genome annotation process (Figure S4)
- The statistics for the different versions of the genome assembly (Table S1)
- The statistics of the annotation (Table S2)

**Figure S1**. Workflow of the genome assembly process.

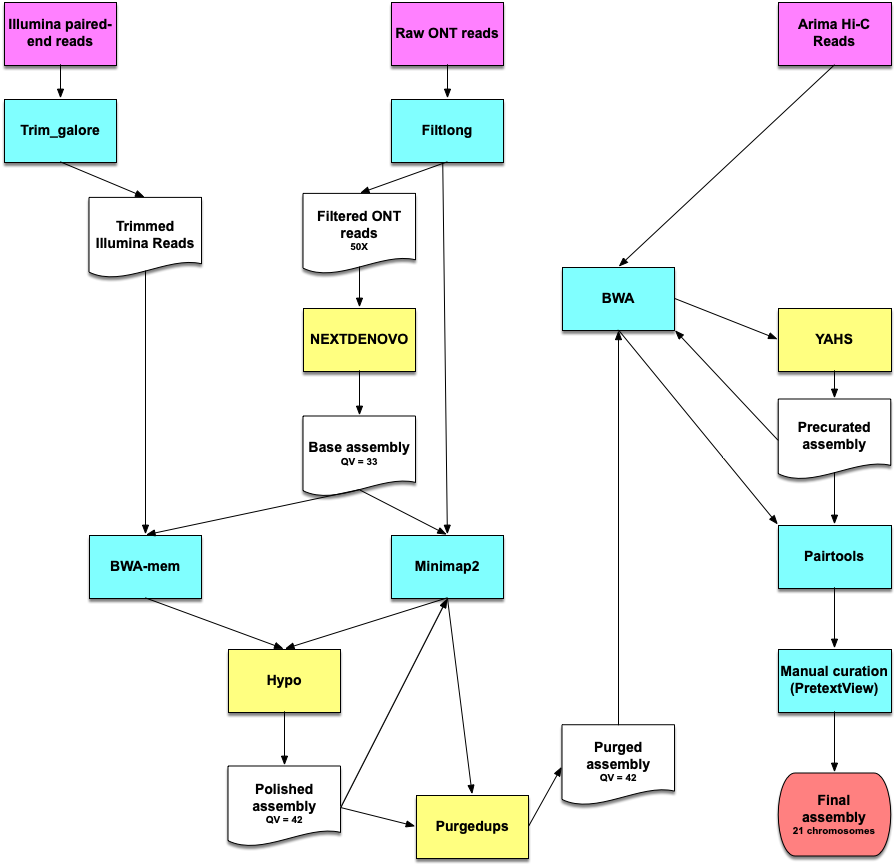

**Figure S2**. Genomescope2 transformed linear plot.

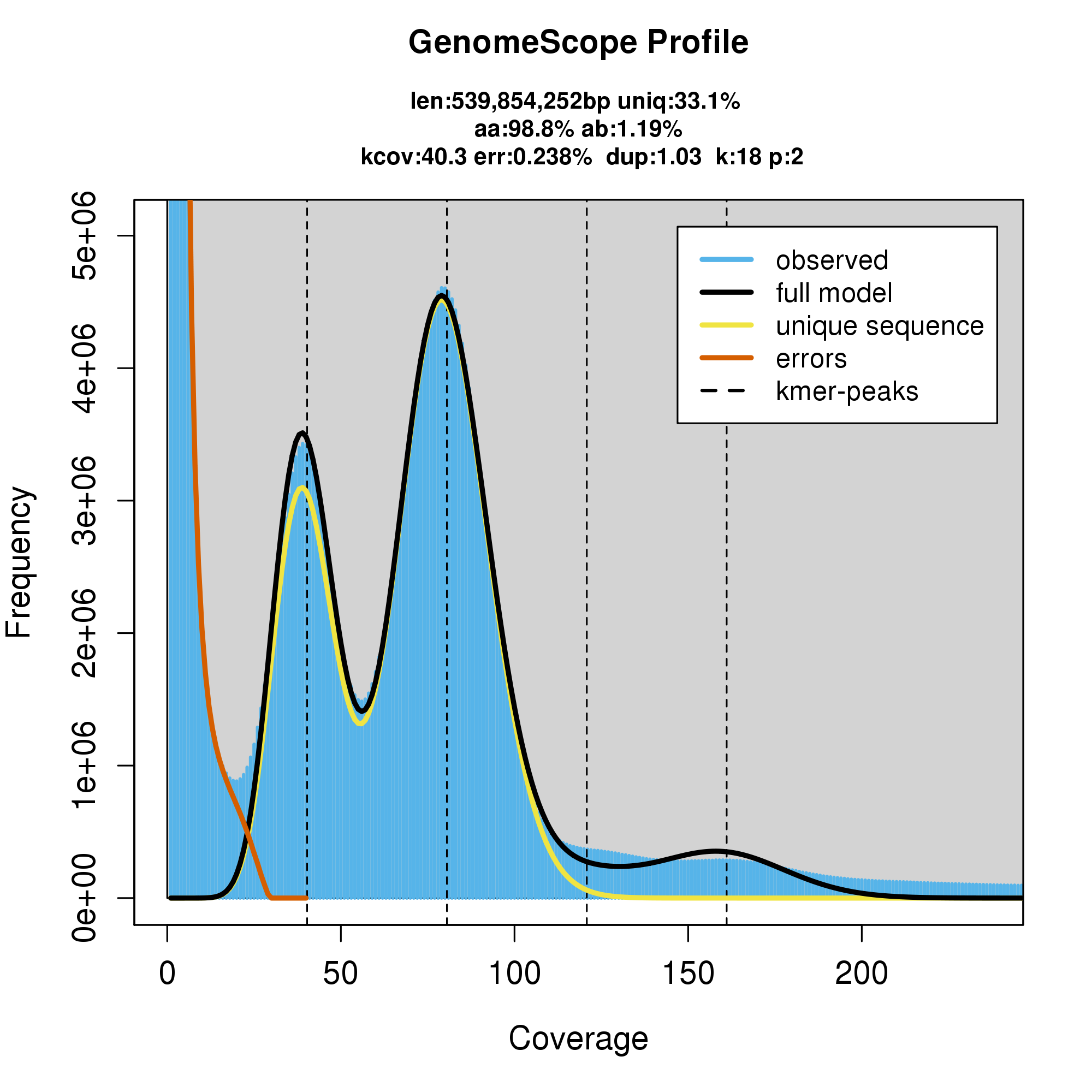

**Figure S3**: **Blob plot of base coverage in illumina against GC proportion for scaffolds in assembly ND_hypo1_purged.** Scaffolds are coloured by phylum. Circles are sized in proportion to scaffold length. Histograms show the distribution of scaffold length sum along each axis.

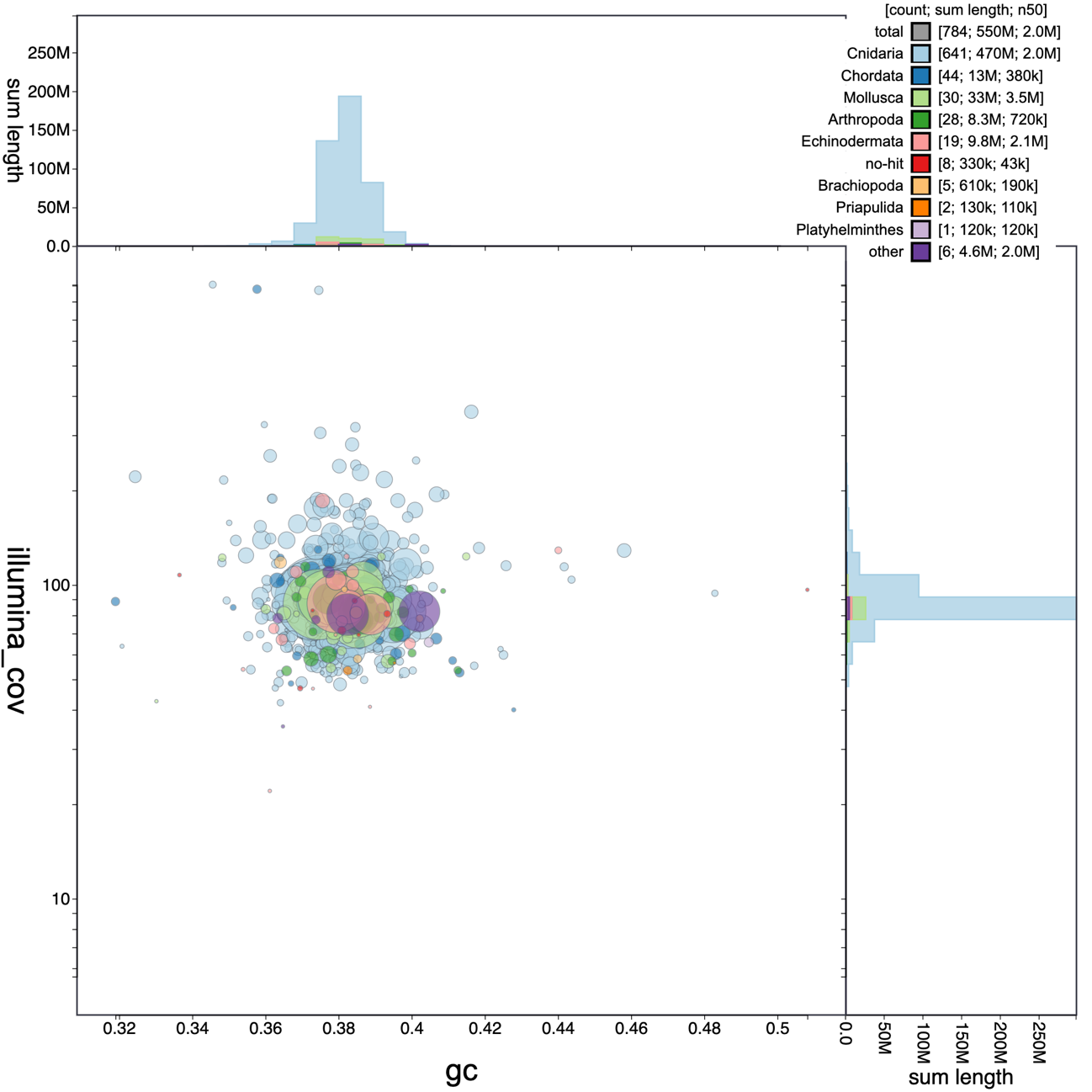

**Figure S4**: workflow of the genome annotation process

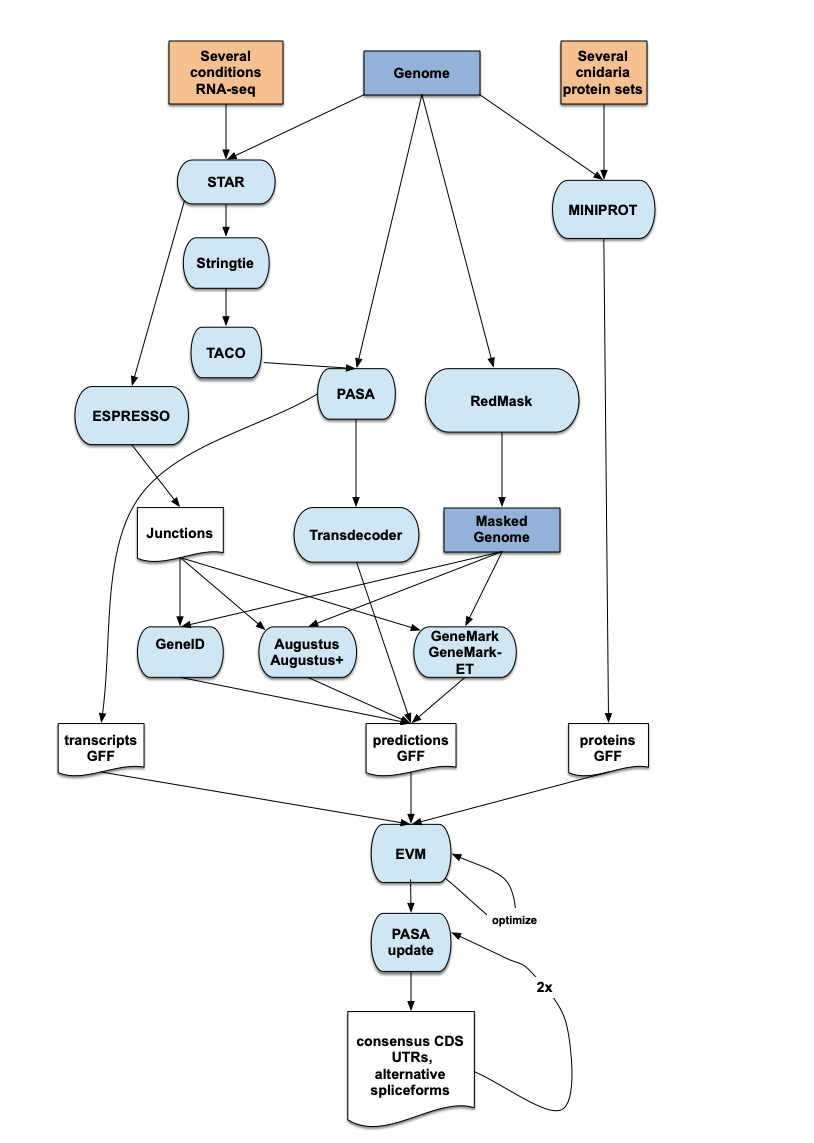

Table S1: Genome assembly statistics

| **Assembly** | **Nextdenovo (ND)** | **ND + hypo** | **ND + hypo + purged** | **jaCorRubr1.1** |
| --- | --- | --- | --- | --- |
| **Contig N50** | 1,993,440 bp | 1,992,814 bp | 2,029,805 bp | 1,637,482 bp |
| **Scaffold N50** | 1,993,440 bp | 1,992,814 bp | 2,029,805 bp | 16,290,029 bp |
| **Scaffold L50** | 84 | 84 | 79 | 8 |
| **Total sequences** | 876 | 876 | 784 | 326 |
| **Assembly span** | 567,713,602 bp | 567,661,090 bp | 545,517,441 bp | 474,689,186 bp |
| **BUSCO* complete** | 84.3% | 88.1% | 88.2% | 74% |
| **BUSCO* duplicated** | 2.3% | 2.5 % | 1.2% | 0.9% |
| **QV** | 33 | 42 | 42 | 42 |
| **K-mer completeness** | 83.8% | 85.8% | 84.9 | 75.2 |

*BUSCO v5 metazoa_odb10 database

Table S2: Genome annotation statistics

|  | **CORRUBR1A annotation** |
| --- | --- |
| Number of protein-coding genes | 39,114 |
| Median gene length (bp) | 1,869 |
| Number of transcripts | 44,624 |
| Number of exons | 180,717 |
| Number of coding exons | 174,919 |
| Median UTR length (bp) | 316 |
| Median intron length (bp) | 780 |
| Exons/transcript | 5.09 |
| Transcripts/gene | 1.14 |
| Multi-exonic transcripts | 63% |
| Gene density (gene/Mb) | 82.40 |
